# The development of an asymmetric accessory olfactory bulb in *Octodon degus* and the influence of social behavior in its establishment

**DOI:** 10.64898/2025.12.30.696934

**Authors:** Pedro Fernández-Aburto, Scarlett E. Delgado, Jaime Mulet, Karina Buldrini, Raúl Sobrero, Jorge Mpodozis

## Abstract

The vomeronasal system (VNS) has been extensively associated with the regulation of sexual and social behaviors in mammals. This system comprises two types of vomeronasal neurons sending parallel projections from the vomeronasal organ (VNO) to the anterior and posterior portions of the accessory olfactory bulb (aAOB and pAOB, respectively). Members of caviomorph rodents develop either a larger aAOB or pAOB bias, which has been associated with ecological traits related to the pheromonal communication displayed by these species. However, it is still unknown how and when such AOB asymmetries emerge and whether experience may influence their establishment. To explore these factors, we studied the development of the VNS in *Octodon degus*, a caviomorph that exhibits an asymmetric AOB, in which the aAOB is larger and has more glomeruli than the pAOB. We found that both VNO and AOB are present at birth, but exhibiting substantial immature traits. Both VNS structures develop postnatally, showing mature patterns by the end of the first postnatal month. By the second postnatal week, aAOB becomes larger in size than pAOB, however, in the glomerular layer such bias appears later, by the end of the first postnatal month. The AOB asymmetry increases in extent until adulthood, as in this period the pAOB shows a minor growing rate than the aAOB. Additionally, we found that animals raised in restricted social contexts displayed a reduced AOB asymmetry in comparison to wild degus. Overall, we suggest that social experiences contribute to the development of the AOB asymmetry in degus.

**Key points:**

- In degus, the vomeronasal organ and the accessory olfactory bulb are present by birth, both displaying immature anatomical traits. More mature, adult-like vomeronasal structures are suggested to be present by the end of the first postnatal month.
- A bias toward a larger volume of the anterior portion of the accessory olfactory bulb compared to its posterior portion, as reported in adult degus, emerges within the first postnatal month, but continues to increase in extent until adulthood.
- The AOB asymmetry is reduced in captivity-reared compared to wild-captured degus and it is absent entirely in individuals raised in isolation conditions.

## 1. Introduction

In mammals, most social and reproductive behaviors are orchestrated via the accessory olfactory or vomeronasal system (VNS), which is mainly activated by pheromone-like semiochemicals (for review see Tirindelli et al. 2009). Molecules reaching the vomeronasal organ (VNO) activate one of two distinct populations of neurons expressing either V1R or V2R metabotropic receptors (Jia and Halpern 1996; Halpern et al. 1998). Additionally, these populations differ significantly in gene expression, stimulus affinity and behavioral modulation. V1R neurons are primarily activated by small, volatile pheromones (Inamura et al. 1999; Inamura and Kashiwayanagi 2000; Leinders-Zufall et al. 2000; Sugai et al. 2006), whereas V2R neurons respond mainly to large, non-volatile pheromones (Leinders-Zufall et al. 2004, 2009; Kimoto et al. 2005). Activation of the V1R pathway is mainly involved in male-female attraction, whereas the V2R pathway is associated with male-male aggression (Dudley and Moss 1999; Inamura et al. 1999; Kumar et al. 1999; Matsuoka et al. 1999; Inamura and Kashiwayanagi 2000; Peele et al. 2003; Sugai et al. 2006; Chamero et al. 2011). From the VNO, V1R and V2R expressing neurons send parallel projections to the anterior and posterior portions of the accessory olfactory bulb (aAOB and pAOB, respectively) (Jia and Halpern 1996; Halpern et al. 1998). In both AOB portions, the arborizations of VNO neurons form synaptic contacts with mitral and tufted neurons in sphere-like structures called glomeruli. Efferent fibers from each glomerulus project to multiple telencephalic regions, especially those involved in neuroendocrine control (Scalia and Winans 1975; Newman and Winans 1980; von Campenhausen and Mori 2000; Martinez-Marcos 2009). Juxtaglomerular and granular interneurons are abundant in both AOB portions, connecting with glomeruli and exerting dynamic modulation over neuronal activity (Jia et al. 1999, Maksimova et al. 2019).

The presence of dichotomous VNS pathways has been widely reported across mammals, from marsupials such as opossums (Halpern et al. 1995) to eutherian species (Berghard and Buck 1996; Villamayor et al. 2018). However, some species within different mammalian clades exhibit non-dichotomous VNS configurations. For example, in the Madagascar tenrec (*Echinops telfairi*), an Afrotheria species, some V1R neurons project their axons into pAOB glomeruli and some V2R neurons project their axons into aAOB glomeruli (Suárez et al. 2009a). The complete loss of the V2R pathway has been extensively reported in members of the superorder Laurasiatheria, as well as in primates (Takigami et al. 2000, 2004; Ngwenya et al. 2011; Kondoh et al. 2017). It has also been observed in the basal Rock Hyrax (*Procavia capensis*, Proboscidea) and the California ground squirrel (*Otospermophilus beecheyi*, Rodentia) (Suárez et al. 2011a). The loss of the V2R pathway has been associated with a reduction in body-to-body contact behaviors that rely on non-volatile cues, such as male-male aggression, especially in species that predominantly rely on visual cues to assess social and/or sexual status, particularly those exhibiting sexual dimorphism. Lastly, the complete absence of both VNS pathways has also been reported in some ape species, in bats, and in some species with a strictly aquatic lifestyle (Meisami and Bhatnagar 1998; Zhao et al. 2011). Overall, while some traits of the VNS are conserved across groups, notable differences among species are evident; most of these variations seem to be related to ecological and behavioral traits specific to each species.

Members of South American caviomorph rodents have been reported to exhibit a remarkable bias toward either aAOB or pAOB having a larger size. For example, *Octodon degus*, a highly social octodontid living in semiarid habitats, possesses an aAOB that is twice the size of the pAOB, with more and larger glomeruli (Suárez and Mpodozis 2009b). In contrast, the capybara (*Hydrochoerus hydrochaeris*), a social rodent inhabiting wet environments, shows the opposite bias, with the pAOB being larger and containing more glomeruli than the aAOB (Suárez et al. 2011b). It was suggested that degus and capybaras would be engaged in contrasting chemosignaling using preferentially volatile and non-volatile cues respectively, as well as in different social interactions involving pheromone sampling (Suárez and Mpodozis 2009b; Suárez and Mpodozis 2011b). Such differences in pheromonal communication would be related to the emergence of a bias towards a larger aAOB or pAOB, as exhibited for caviomorph species.

Recently, we proposed that variations in social behavior could be a driving factor behind AOB asymmetry in closely related species. We examined the AOB morphometry of *Octodon lunatus*, a sister species of *O. degus*. Compared to the highly social degus, *O. lunatus* exhibits a lower level of social interaction (Sobrero et al. 2014). Like degus, *O. lunatus* has a larger overall aAOB, but they display a smaller glomerular layer, similar in volume to the pAOB (Fernández-Aburto et al. 2019). Thus, AOB asymmetry develops differently among closely related species that exhibit contrasting social behavior. This cumulative evidence suggests that natural history traits, such as social behavior, may serve as predictors of the AOB shape in caviomorph rodents.

The goal of this study was to understand how and when the AOB asymmetry is established and how social behavior influences its development in *O. degus*. Degus are well-studied organisms in terms of their development and ecology. They are diurnal rodents characterized by a long gestation period of approximately 3 months (Rojas et al. 1982). Typically, females produce around six precocial pups with fur and open eyes. In the wild, pups begin scavenging around their first week, but parental care is extended for the end of the first month (Veloso 1997; Reynolds and Wright 1979). Both males and females reach sexual maturity at approximately 3.5 months, with the first breeding usually occurring near the six-month age (Lee 2004; Hummer et al. 2007; Mahoney et al. 2011). In the wild, degus form complex social structures. During the reproductive season, males monopolize groups of 5-6 females (Fulk 1976). A vast diversity of social behaviors including communal burrowing (Ebensperger and Bozinovic 2000), male dispersion (Quirici et al. 2011), communal nursing (Ebensperger et al. 2004; Jesseau et al. 2009), vigilance for predators (Ebensperger et al. 2006), social foraging (Vasquez 1997), and scent marking of territory using olfactory cues (Ebensperger and Caiozzi 2002) are commonly displayed. The prolonged juvenile period and intricate social interactions in which pheromone signaling may act as a mediator, make degus an interesting species for studying the VNS ontogeny and the influence of social experience in its development.

In this study, the overarching question is whether the development of the AOB asymmetry in degus is influenced by social experience. Based on the evidence pointing to a role of social experience on AOB shaping in octodontids, we tested the hypothesis that the emergence and establishment of the AOB asymmetry in degus occurs during the postnatal period, driven by social experience. First, we studied the anatomical and molecular development of VNO and AOB, to examine the course of maturation of the VNS. Next, by characterizing the morphometry of aAOB and pAOB at different ages, we assessed when the AOB asymmetry (aAOB > pAOB) first appears and how it progresses over time. Finally, we studied the influence of social context on the establishment of the AOB asymmetry. We report here that in degus, the transition to a mature, adult-like VNS occurs during the first postnatal month. A larger aAOB than pAOB emerges within the first postnatal month, however, the asymmetry remains growing over later ages, reaching a maximum extent by adulthood. Finally, we found that degus raised in captivity and controlled isolation conditions developed less asymmetric and symmetric AOBs, respectively. Our findings support the idea that social experience plays a key role in shaping the AOB in degus.

## 2. Methods

### 2.1 Animals

All procedures were performed in accordance with guidelines established by the University of Chile Bioethical Committee (120731-DB-FSH-UCH) and adhered to Chilean laws for animal captures [permits 1-130 154.2010 (7989), 1-109.2011 (6749), 1-90.2011 (4731) and 1-95-2012 (4486) by the Servicio Agrícola y Ganadero and 013/2011 by the Corporación Nacional Forestal].

Captivity-reared degus: 6 pregnant *O. degus* were captured in the wild at the end of the winter season (August-September) and housed in the animal facility at the University of Chile. They were kept in groups of 2-3 individuals, with water and food *adlibitum*. Litters from one individual were obtained before parturition, at approximately 60 and 80 days of gestation (G60-G80) (see *tissue preparation*). The age of each fetus was estimated based on embryological data from Rojas et al. 1982. In captivity, each mother gave birth to 5-7 pups. The day after birth, both the mother and her offspring were moved to a different cage, and they remained together until weaning. The VNO and olfactory bulbs of 5-7 individuals raised in captivity were collected at different postnatal days: 1day (P1), 15 days (P15), 30 days (P30), 60 days (P60), 150 days (P150) and adults (greater than 180 days). Adults were defined as individuals in the breeding stage, typically between 6 months and 1 year (Hummer et al. 2007).

Field-captured degus: The olfactory bulbs of adult, field-captured *O. degus*, were obtained from parallel experiments conducted in our laboratory. For comparative purposes, we also collected olfactory bulbs of adults, field-captured *Octodon lunatus* and adults, captive-reared rats (*Rattus norvegicus*).

Isolation-reared degus: For isolation experiments, six degus of one month of age (P30) were separated from the others and housed in pairs within specially designed chambers (50 x 60 x 40 cm) that prevented direct contact with other individuals and their odors, allowing them to grow until reaching adulthood. Animals were kept with water and food *adlibitum.* Due to methodological reasons, only the olfactory bulbs of three isolated-raised animals were suitable for analysis.

### 2.2 Tissue preparation

All animals were deeply anesthetized with ketamine and xylazine (2.4 mL/kg and 1.2 mL/kg respectively) and transcardially perfused with 0.9% saline solution, followed by 4% paraformaldehyde in 0.1M PBS. The brains were then removed, weighed, and post-fixed until they underwent further processing. The olfactory bulbs were dissected and placed in a 30% sucrose solution in 0.01 M PBS for cryoprotection until they sank. Sagittal sections of 40μm thickness were obtained using a freezing microtome (Leitz Wetzlar 1400, Leitz, Germany). The sections were collected into two series at 40μm intervals and stored in a solution of 0.01 M PBS with sodium azide for subsequent processing. VNO samples were surgically extracted from each individual’s nose and embedded, either in gelatin (porcine skin gelatin, Sigma-Aldrich) or paraffin (Paraplast ®). From gelatin-embedded samples, 30μm transverse sections of the VNO were obtained using a freezing microtome, processed for immunohistochemistry and counterstained with cresyl violet (Certistain®). Paraffin-embedded samples yielded 10μm transverse sections, which were obtained using a paraffin-slicing microtome (Leica Biosystems), mounted on silane-coated slides, and stained with haematoxylin-eosin solution and OMP immunostaining.

### 2.3 Immunohistochemistry

Sagittal sections of the olfactory bulbs were processed as free-floating sections. They were incubated with 3% hydrogen peroxide (H_2_O_2_) and 10% methanol for 10 min, followed by incubation in sodium citrate buffer (10 mM, pH 6.0) at 80 ^◦^C for 30 min. The sections were then blocked with 5% normal goat serum (NGS) in 0.05% Triton X-100 in PBS (PBS-T) for 1 h. Subsequently, the slices were incubated with one of the following primary antibodies: goat anti-G_αi2_ mouse, monoclonal 1:500, cat no. sc-13534, Santa Cruz Biotechnology), anti-OMP, mouse monoclonal (1:500, cat no. sc-365818, Santa Cruz Biotechnology) and anti-VGLUT2, rat monoclonal (1:1000, clone N29/29, NeuroMab), all diluted in 3% NGS in PBS-T, at 4^◦^C for 2 h. After rinsing, sections were incubated with one of the following secondary antibodies: biotinylated anti-mouse made in goat (1:500, cat no. BA-9200, Vector Laboratories) or rhodamine-coupled anti-mouse made in goat (1:500, cat no. 31660, Invitrogen) for 2 h. The biotinylated sections were then treated with the avidin-biotin complex (ABC Elite Kit; Vector Laboratories) for 1hour, followed by a reaction with 0.25 mg/ml of 3,3-diaminobenzidine (cat no. D-5905, Sigma-Aldrich) in PBS with 0.001% H_2_O_2_ for 3 min. The DAB-stained sections were mounted and counterstained with either cresyl violet or GIEMSA. The fluorophore-labeled sections were washed with PBS and counterstained with DAPI for 5 minutes, then mounted and coverslipped with Fluosave™ Reagent (Merck Millipore). For gelatin-embedded VNO samples, sections were mounted on silane-coated slides, immunoreacted with anti-G_αi2_ using a modified immunohistochemistry protocol for mounted sections, and then reacted with DAB.

### 2.4 Stereology

Volume measurements of both AOB portions and their corresponding glomerular layers (GL) were estimated using a stereological method. The method involved the measurement of cross section areas of each AOB structure, in cresyl-violet stained serial sections separated by 40μm. Stereological measurements were performed using Stereo Investigator 8.0 software (MBF Bioscience, Williston, Vt., USA), with a CCD video camera (Microfare, Optometrics ®) coupled to a Nikon Eclipse E400 bright-field microscope. All sampling schemes were optimized based on standard strategies for stereological analysis. Using the Cavalieri stereological estimator (CE), we measured the volume of the overall AOB subdomains and their corresponding GLs, employing a 30μm square grid at ×40 magnification for each sample. The Gundersen coefficients of error (GCE), used to estimate the precision of each volume measurement, were calculated for each sample (m=1). GCE values ranged between 0.002 and 0.008 for the aAOB (P1: 0.003-0.005; P15: 0.003-0.005; P30: 0.002-0.004; P60: 0.003-0.007; P150: 0.003-0.008 and adults: 0.002-0.007) and between 0.003 and 0.007 for the pAOB (P1: 0.004-0.006; P15: 0.004-0.015; P30: 0.002-0.005; P60: 0.003-0.006; P150: 0.004-0.005 and adults: 0.003-0.009). For comparative purposes, we divided the estimated volume of a given structure by brain volume, considering any allometric effects. Brain volume was calculated by dividing the brain mass by fixed brain tissue density (1.036g cm^−3^; Barber et al. 1970). Thus, volumes are expressed as relative to the brain volume. For AOB and GL ratio estimations, we also included ratio values obtained from animals for which the size of these structures were not normalized for brain volume.

### 2.5 Statistical Analyses

Parametric and non-parametric statistical tests were applied based on the results of the Shapiro-Wilk test for normality. A one-way ANOVA or Kruskal-Wallis test was employed to compare the AOB and GL volumes among age groups and among adult degus raised in different contexts. A paired *t*-test or Wilcoxon-matched pairs test was used to compare aAOB and pAOB volumes within each age group. All analyses were performed using SigmaPlot 14.0. Differences with *p*-values less than 0.05 and 0.01 were considered statistically significant.

### 2.6 Figures processing

Bright-field photomicrographs were captured using an Olympus microscope coupled with a SPOT advanced camera (Diagnostic Instrument). Fluorescent images were obtained with an Olympus BX-60 microscope equipped with a spinning disk confocal scanner unit and a Hamamatsu C10600 camera. Fluorescent images were acquired using CellSense® software and processed with Adobe Photoshop (Adobe Systems, San Jose, CA). Contrast and brightness adjustments were made only when necessary. All graphs were generated using custom scripts written in MATLAB (Mathworks, Inc). The intensity of OMP immunoreactivity was measured from VNO and AOB images using ImageJ.

## 3. Results

### 3.1. In degus, VNO and AOB are present before birth, but exhibit important immature features

First, we evaluated the development of VNS structures in degus, focusing on the timing of their emergence and their subsequent anatomical and molecular maturation. Samples from VNO and AOB from individuals at prenatal stages between 60 and 80 days of gestation (G60, G80), were analyzed. In degus, the VNO and AOB were clearly observable near the end of gestation (Figure 1A, B). At G80, all of the AOB layers previously described in adults (Suárez and Mpodozis 2009b) were distinguishable: vomeronasal nerve layer (VNL), glomerular layer (GL), external plexiform layer (EPL), mitral tufted cell layer (MT/L), lateral olfactory tract (lot) and granular layer (GrL) (Figure 1C). However, notably immature traits were also evident during this period. The GL was thin, containing few glomeruli, many of which were surrounded by very few interneurons. A majority of neuronal cell bodies were located primarily at the base of the GL. At this stage, somata of mitral and tufted neurons formed a thick layer, with some scattered across the EPL. Consistent with our previous report on adult degus (Suárez and Mpodozis 2009b), the AOB of G80 individuals showed anti-G_αi2_ immunoreactivity (G_αi2_-ir) restricted to the VNL and GL of the aAOB (Figure 1D). Moreover, the G_αi2_-ir pattern coincided with the presence of an indentation that separates the two AOB portions, which is more clearly observed at the MT/L (Figure 1B, C and D). These findings suggest that axons from V1R and V2R neurons in *O. degus* project separately and reach the anterior and posterior portions, respectively, of AOB before birth.

**Figure 1.**
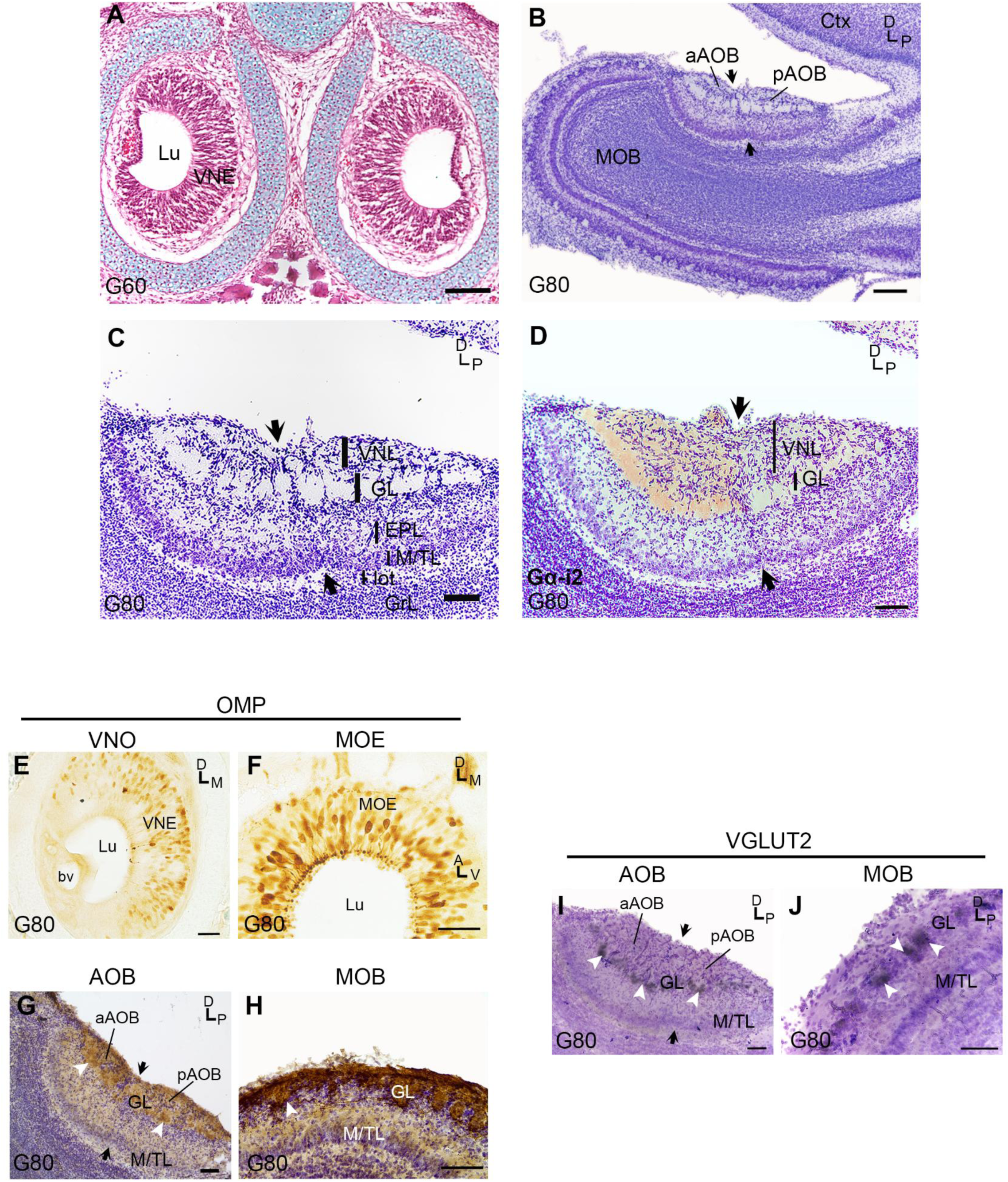
VNS of *O. degus* at middle/late prenatal period. A) Transverse section of the nose at 60 gestation days (G60), showing both left and right VNO. Section was stained with hematoxylin/eosin and Alcian Blue (cartilage, blue). Lu, lumen; VNE, vomeronasal epithelium. Scale bar 100 µm. B) Sagittal section of the olfactory bulb at G80, the AOB is in its dorsal-caudal region. The separation between aAOB and pAOB is indicated with black arrows. Section was counterstained with cresyl violet. Ctx, frontal cortex; MOB, main olfactory bulb. Scale bar 200 µm. C) Magnified view of the AOB at G80, showing its multilayered organization. VNL, vomeronasal nerve layer; GL, glomerular layer; EPL, external plexiform layer; M/TL, mitral/tufted cells layer; lot, lateral olfactory tract; GrL, granular layer. Bar, 100 µm. D) G_αi2_-ir, located exclusively in the aAOB (VNL+GL). Section was counterstained with cresyl violet. Bar, 100 µm. E-J) Olfactory and vomeronasal neurons show limited expression of key olfactory markers by G80. E, F) Presence of sparce vomeronasal neurons exhibiting OMP-ir in the VNE (E) and MOE (F). Lu, lumen; VNE, vomeronasal epithelium; MOE, main olfactory epithelium. Scale bar 100 µm. G, H) OMP-ir observed in the aAOB and pAOB (G) and MOB (H) (VNL+GL). Glomeruli profiles at AOB are indicated with arrowheads. Scale bar 100 µm. I, J) VGLUT2-ir glomeruli (dark stain, white arrowheads) present at GL of both, AOB (I) and MOB (J). Sections were counterstained with GIEMSA. The separation between aAOB and pAOB is indicated with black arrows. Glomeruli profiles are indicated with arrowheads. Scale bar 100 µm.

To assess the maturation state of VNS neurons projecting to the AOB at prenatal ages, we evaluated the expression of two molecular markers associated with mature olfactory neurons: the olfactory marker protein (OMP) and the vesicular glutamate transporter 2 (VGLUT2). OMP is reported to be exclusively expressed in differentiated, functional olfactory neurons in mice (Monti-Graziadei et al. 1977; Lee et al. 2011), rats (Farbman and Margolis 1980) and primates (Dennis et al. 2004). In G80 degus, we observed scattered OMP-immunoreactive (OMP-ir) neurons throughout the vomeronasal epithelium (VNE) (Figure 1E). OMP-ir was detected in dendrites, somatas and axon bundles. Unlike in the VNO, a large number of OMP-ir neurons were present in the main olfactory epithelium (MOE) at this stage (Figure 1F). In the AOB, OMP-ir was restricted to the VNL and GL in both AOB portions (Figure 1G), consistent with adult patterns. A similar distribution was observed in the main olfactory bulb (MOB) (Figure 1H). Regarding VGLUT2, its distribution is restricted to the synaptic vesicles of glutamatergic olfactory-type neurons (Herzog et al. 2001; Nakamura et al. 2005). In G80 degus, VGLUT2-ir was confined to the glomeruli of both AOB (Figure 1I) and MOB (Figure 1J), similar to what was found in adults. Overall, despite the presence of both VNO and AOB structures before birth, immature anatomical and molecular traits are still observed.

### 3.2 Mature anatomical traits in the AOB are first established during the early postnatal time

Next, we examined the anatomy of the olfactory bulb across the postnatal development. The MOB exhibited glomeruli well surrounded by interneurons at P1 (Figure 2A). At this stage, however, glomeruli were small, and in some regions, they were not organized in a row, a pattern commonly observed in adults. Glomeruli of MOB increased in size and acquired a more organized pattern across later postnatal development (Figure 2B-E).

**Figure 2.**
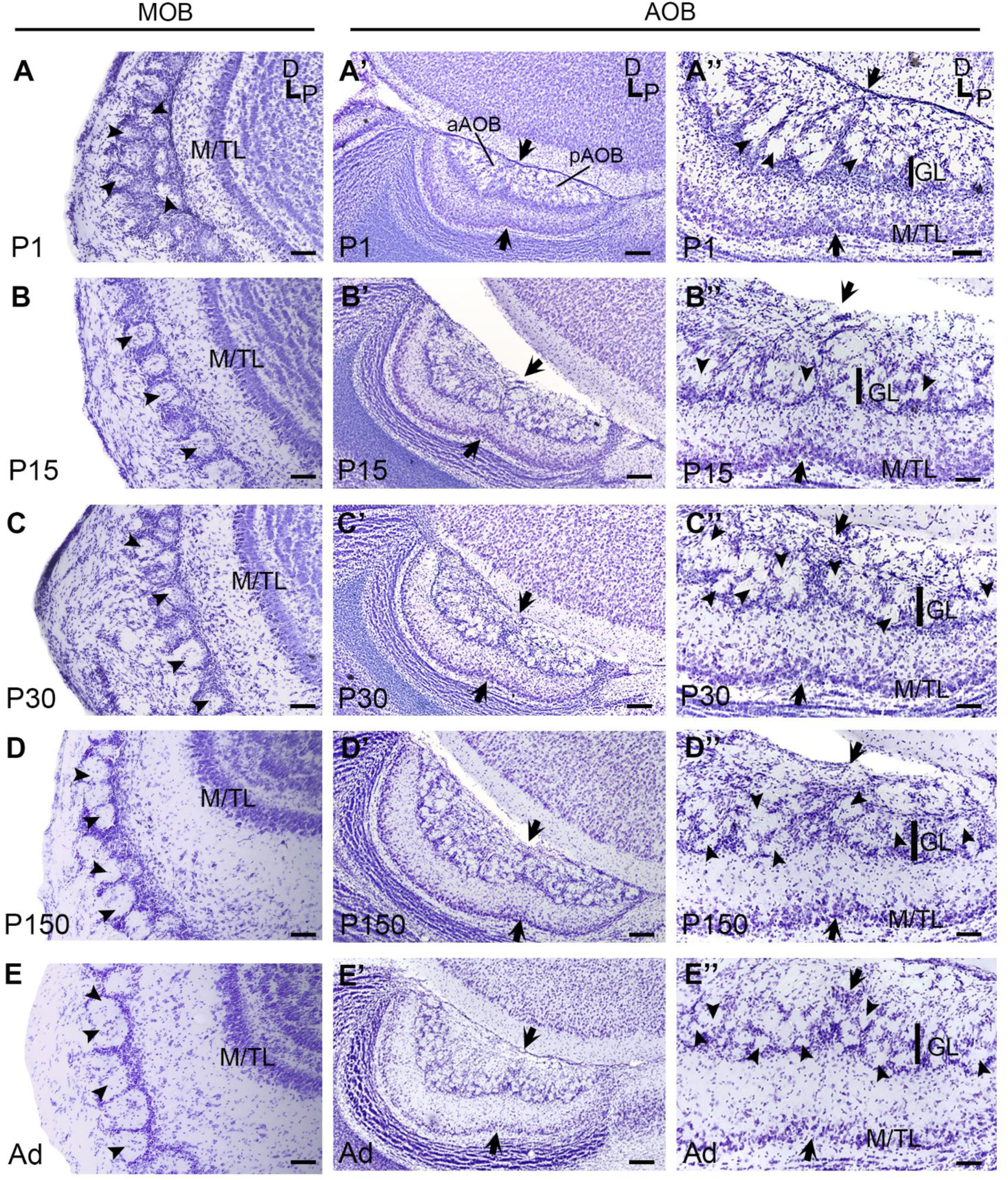
Anatomy of olfactory bulbs of *O. degus* across postnatal development. A-E) Sagittal sections showing the rostral-ventral portion of MOB at P1(A), P15(B), P30 (C), P150 (D) and adults (E). Glomeruli profiles at AOB are indicated with white arrowheads. Scale bar, 200 µm. A’-E’) Sagittal sections showing the AOB at P1(A’), P15(B’), P30 (C’), P150 (D’) and adults (E’). Scale bar, 200 µm. A’’-E’’) Magnified view showing layers of both AOB portions at P1(A’’), P15(B’’), P30 (C’’), P150 (D’’) and adults (E’’). Sections were stained with cresyl violet. GL, glomerular layer; M/TL, mitral/tufted cells layer. The separation between aAOB and pAOB is indicated with black arrows. Glomeruli profiles at AOB are indicated with arrowheads. Scale bar, 100 µm.

In comparison to G80, AOB of P1 degus the GL increased in thickness, and few well-defined glomeruli surrounded by interneurons were visible (Figure 2A’, A’’). Nonetheless, large regions of the aAOB still showed glomeruli poorly surrounded by interneurons, similar to the prenatal period. Additionally, numerous neuronal somata remained at the base of the GL in both AOB portions, similar to the prenatal stages. By P15 (Figure 2B’, B’’), the GL and EPL showed a significant increase in thickness, along with a reduction in the number of neurons at the base of the GL. However, large areas of the GL still exhibited few interneurons. From P30 until adulthood, AOB layers exhibited an increase in thickness, with more pronounced changes observed in the aAOB than pAOB (Figure 2C’, D’, E’). The increase in aGL thickness was associated with larger and numerous sets of glomeruli surrounded by interneurons. In contrast, pGL developed fewer and smaller glomeruli over the same period. Furthermore, the density of juxtaglomerular interneurons within the GL increased over time, correlating with a gradual decrease in the number of cells at the GL base. M/TL also showed a reduction in thickness across development, with mitral and tufted somas becoming more loosely scattered in adults (Figure 2C’’, D’’, E’’). Thus, in comparison to MOB, AOB exhibits a delayed development. Our findings suggest that an adult-like anatomical maturation in the AOB is attained by the end of the first postnatal month.

### 3.3 The expression of key molecular markers in the VNS reaches maturity within the first postnatal month

To explore the maturation of the vomeronasal neurons during development, we evaluated the presence and distribution of key molecular markers at different postnatal ages. In the VNO, the presence of G_αi2_ was restricted to the apical portion of the VNE by P15 (Figure 3A) and persisted into adulthood (Figure 3B). Consistent with prenatal observations, G_αi2_-ir was observed exclusively in aAOB at all the postnatal ages examined (Figure 3C, D, E, F). Likewise, VGLUT2-ir was restricted to the GL of both the aAOB and pAOB from birth until adulthood (Figure 3G, H, I, J, K). Thus, some of the molecular markers expressed by vomeronasal neurons display adult-like distributions by birth.

**Figure 3.**
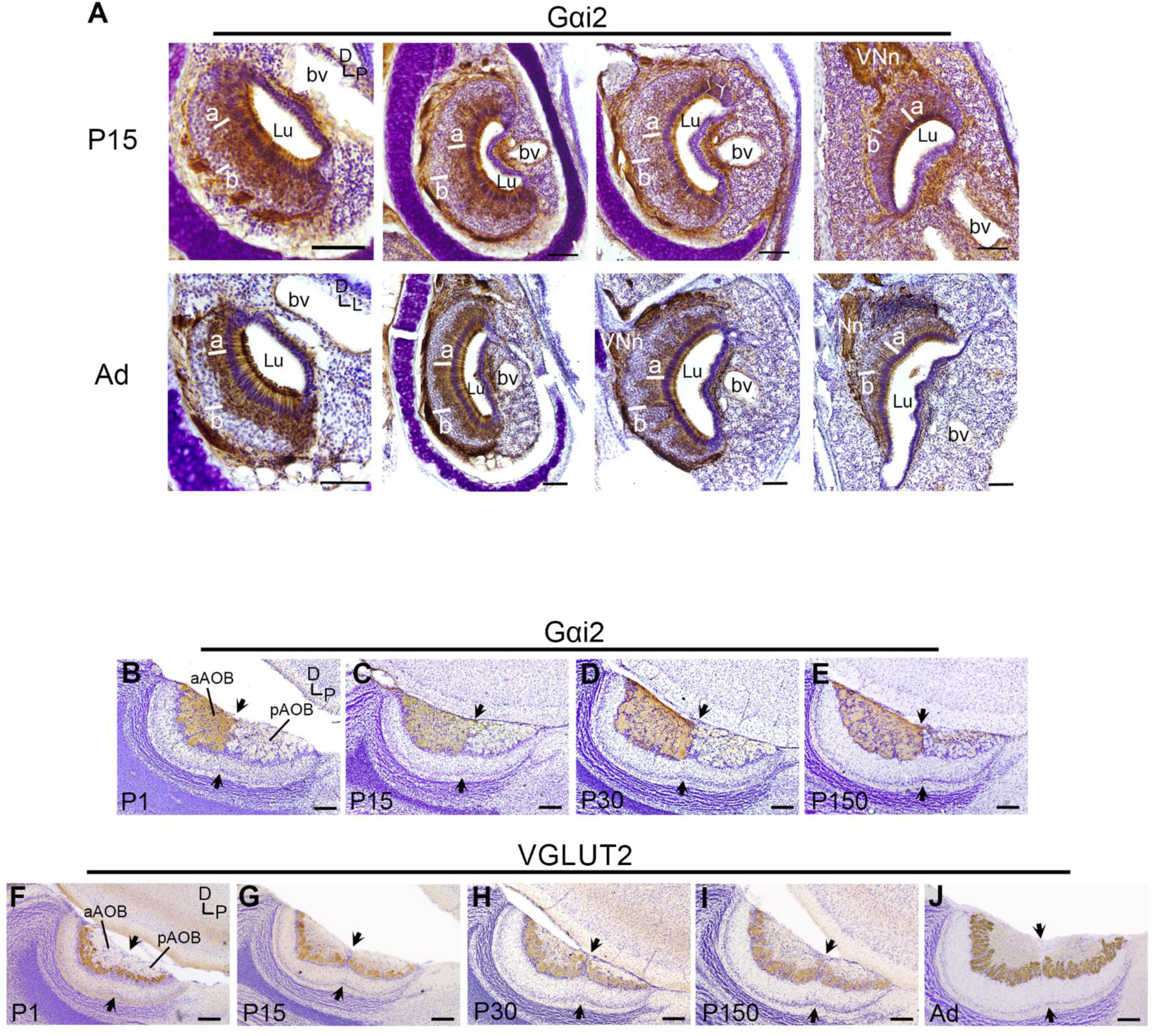
Somas of V1R and V2R neurons and their projections are separated at VNO and AOB from early postnatal ages. A) Transversal sections of P15 (*upper row*) and adults (*lower row*) of VNO, immunostained with G_αi2_-ir. The microphotographs show G_αi2_-ir in different VNO locations in the anterior-posterior axis. Lu, lumen; a, apical; b, basal. Scale bar, 50 µm. B-E) G_αi2_-ir present at VNL and GL of the aAOB; P1 (B), P15 (C), P30 (D), P150 (E). Scale bar, 200 µm. F-J) VGLUT2-ir at GL of the AOB: P1(F), P15 (G), P30 (H), P150 (I) and adults (J). Sections were counterstained with cresyl violet. Scale bar, 200 µm.

Similarly, we assessed the OMP distribution in V1R and V2R neurons across postnatal development. At P1, most neurons in the VNE exhibited OMP-ir (Figure 4A), which contrasted with the scattered pattern OMP-ir observed at G80. OMP expression was present in dendrites, somatas, and nerve bundles surrounding the VNO. By P15, OMP-ir was more intense in neurons located in the most apical portion of the VNE than at its base (Figure 4B). This asymmetric pattern in OMP-ir persisted at later ages evaluated (Figure 4C, D, E). The OMP pattern observed in the apical VNE resembles that of G_αi2_-ir (Figure 3A, B), suggesting that neurons showing strong OMP -ir at VNO are V1R neurons.

**Figure 4.**
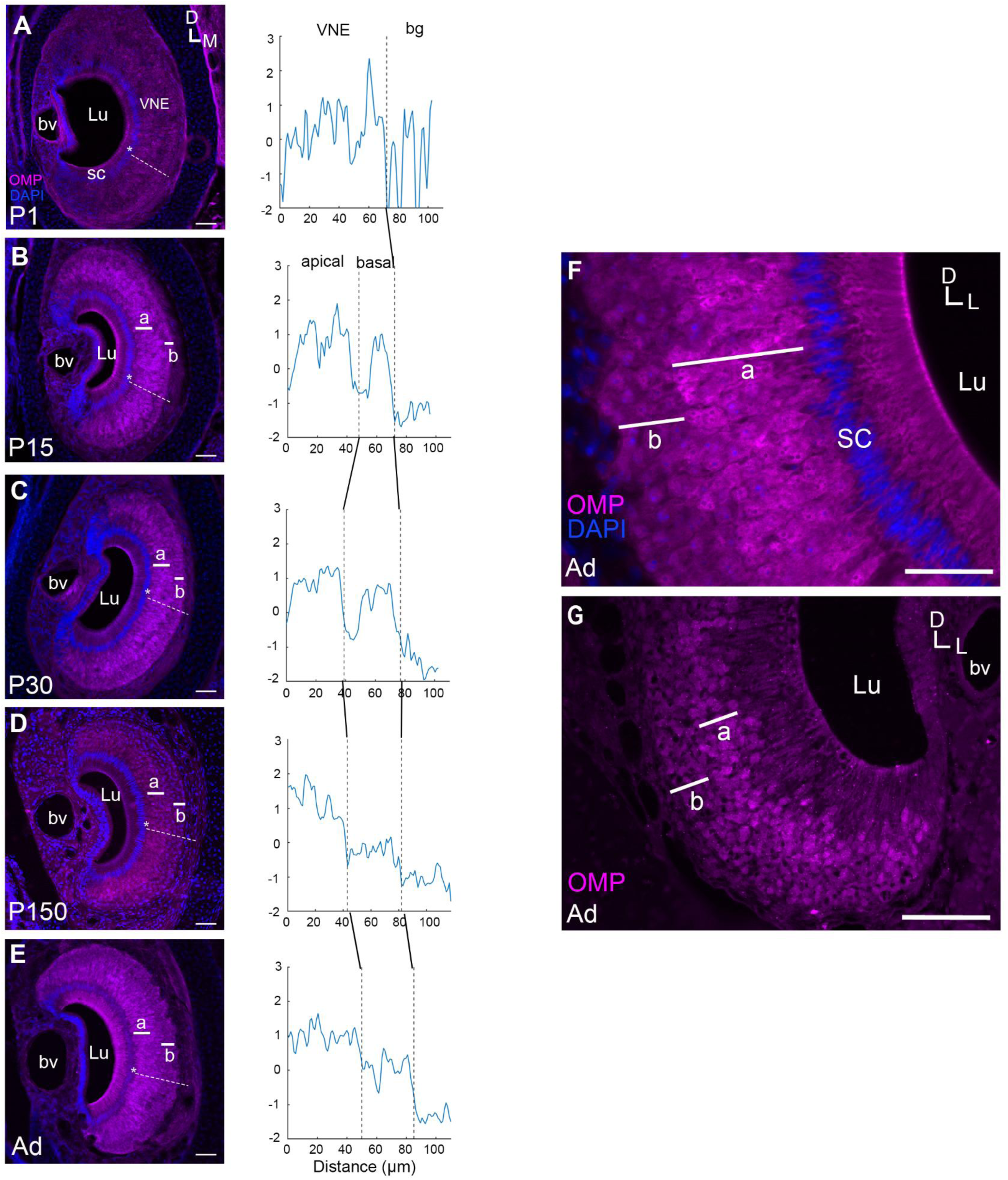
OMP distribution in the VNO across development. A-E) V1R and V2R neurons at VNO exhibit differences in OMP-ir from P15. A) Vomeronasal neurons at P1 exhibit a low, homogeneous OMP-ir in VNO. B) VNO showing a band of apical neurons (a) with a more intense OMP-ir than neurons at the base (b) of the VNE at P15. The difference in OMP-ir persists at P30 (C), P150 (D) and adults (E). Scale bar, 50 µm. White dashed line in all images represents the location of the VNO from which OMP-ir intensity measurements were obtained. Asterisks represent the initial point of the measurement. Graphs showing the OMP-ir intensity measurements for each age are displayed (*right column*). bg, background. F-G). V1R neurons exhibit more intense OMP-ir than V2R neurons at single-cell levels. F) High-magnification microphotographs of the VNE of an adult degu. Apical VNO neurons (a) exhibit more intense OMP-ir than neurons at the base (b) of the VNE, distributed in a 10µm focal plane. Lu, lumen; SC, supporting cells. Scale bar, 50 µm. G) Thin transversal section of VNO obtained from paraffin-embedded samples, showing a stronger OMP-ir at apical VNO neurons than neurons at the base (b) of the VNE. Scale bar, 50 µm.

We examined whether the differences in OMP-ir observed between apical and basal vomeronasal neurons at VNO are present at the single-neuron level. In 10µm thick confocal images of transverse sections of the VNO of adult degus, individual somata in the apical VNE displayed more intense OMP-ir than those at the base (Figure 4F). The same pattern was observed in 10µm transverse sections of the VNO from paraffin-embedded samples (Figure 4G). Thus, the difference in OMP-ir between apical and basal VNO neurons involves differences in OMP expression at the single-neuron level.

Similarly, in the AOB at P1, no significant difference in OMP-ir was found between the aAOB and pAOB (Figure 5A). However, at P15, the GL and VNL at aAOB showed stronger OMP-ir than the pAOB (Figure 5B). By this age, some glomeruli exhibited strong OMP-ir in both AOB portions, most notably in the aGL. The asymmetry in OMP-ir pattern between aAOB and pAOB persisted at later ages (Figure 5C, D). In adults, OMP-ir was uniformly distributed in the VNL and GL in each AOB portion, with the aAOB showing stronger OMP-ir than the pAOB (Figure 5E). To quantify the fluctuations in OMP-ir between aAOB and pAOB across the postnatal development, the OMP-ir ratio (aAOB/pAOB) was calculated at the different ages. A significant increase in the OMP-ir ratio was observed between P1 and the later ages evaluated (one-way ANOVA, F = 8.9, *p <* 0.003, Dunn’s post hoc test, *p* < 0.05; Figure 5F), No differences were observed from P15 ahead (Dunn’s post hoc test, *p* > 0.05). Thus, there is a transition from a uniform distribution of OMP-ir between aAOB and pAOB at birth to a higher OMP presence in aAOB than pAOB by P15, which persist until adulthood.

**Figure 5.**
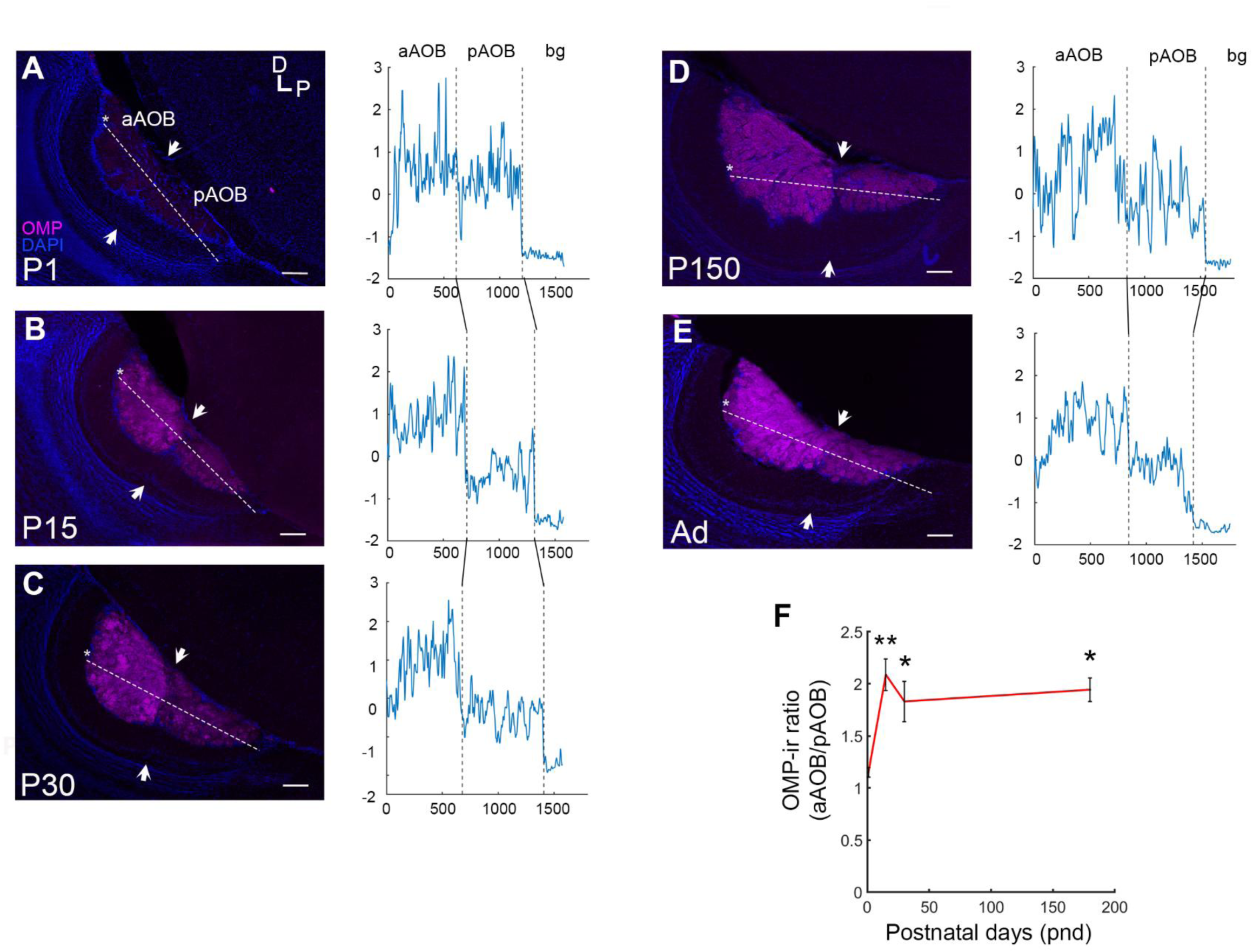
OMP distribution in the AOB across development. A) aAOB and pAOB exhibit similar OMP-ir at P1. B) aAOB exhibits a stronger OMP-ir than pAOB at P15. A-E) The same bias in OMP-ir at AOB persists at P30 (C), P150 (D) and adults (E). Sections counterstained with DAPI. Scale bar, 100 µm. F) OMP-ir ratio (aAOB/pAOB). Graph showing the progression in the OMP-ratio extent through development. Significant differences across ages are shown with asterisk (* p<0.05, ** p<0.01, *** p<0.001).

Overall, our results indicate that the expression pattern of some olfactory markers, such as G_αi2_ and VGLUT2, are established in the VNS by birth. However, others such as OMP exhibit a mature pattern later, starting by the second postnatal week. This suggests that V1R and V2R neurons in degus undergo substantial maturation in the expression of key olfactory markers during the first postnatal month.

### 3.4 The asymmetric expression of OMP in the AOB is also present in a sister species of degus, but not in murids

To examine differences between V1R and V2R neurons within octodontids, we observed the OMP expression in another *Octodon* species. Specifically, we analyzed the OMP-ir in *Octodon lunatus,* an *O. degus* sister species. Lunatus possesses an asymmetric AOB, but unlike degus, a bias in the size of GL between AOB portions is absent (Fernández-Aburto et al. 2019). We found that, similar to wild degus (Figure 6A), field-captured *O. lunatus* exhibited stronger OMP-ir in the VNL and GL of the aAOB compared to the pAOB (Figure 6B). In contrast, in captivity-raised rats (a murid species), the OMP-ir did not show differences between aAOB and pAOB (Figure 6C). Whether the OMP difference between V1 and V2 neurons is a conserved trait within the *Octodon* species deserves further investigation.

**Figure 6.**
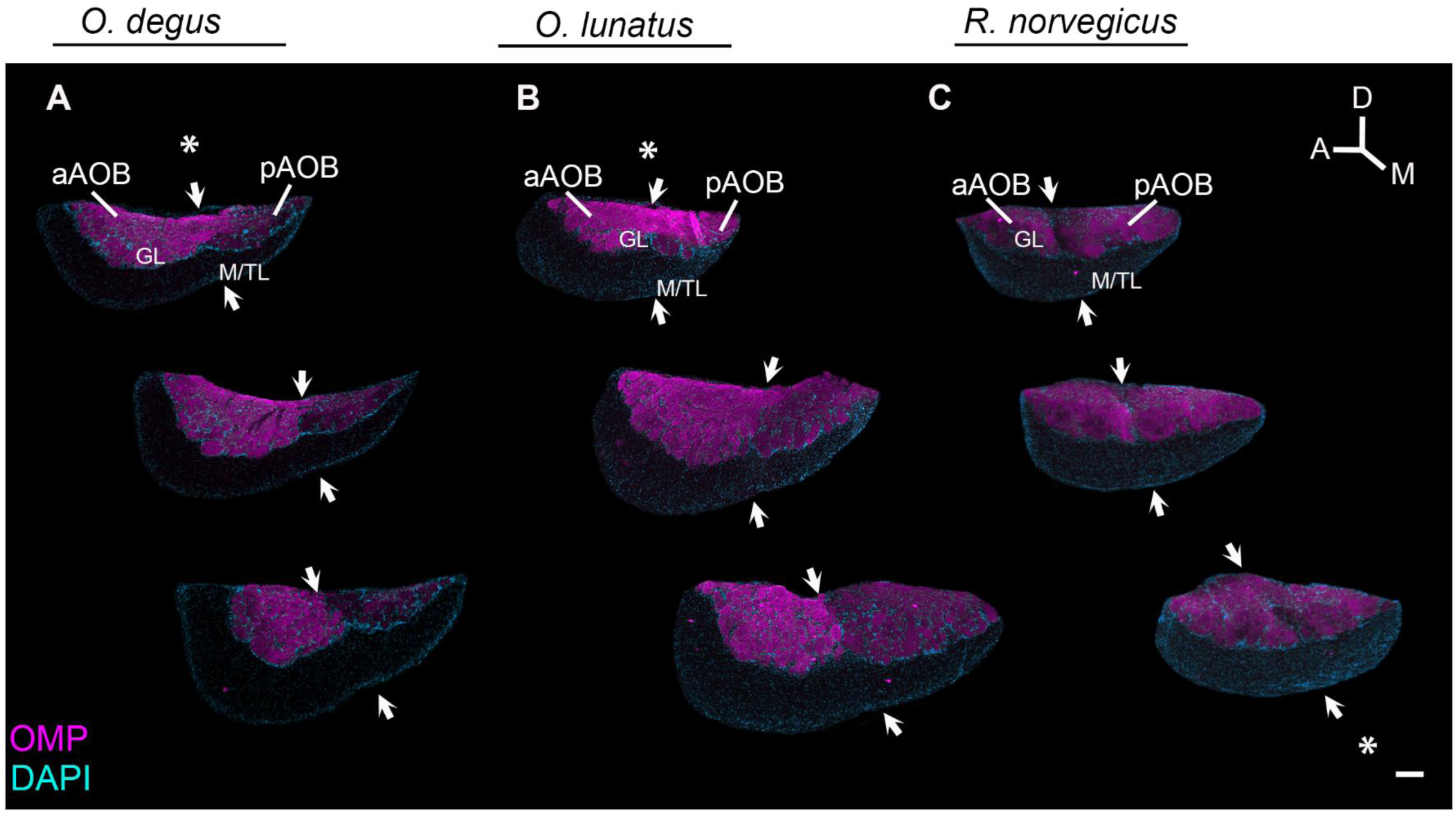
A more intense OMP-ir at aAOB than pAOB is also found in *O. lunatus*, but not in murids. A, B) AOB sections ordered from lateral (top) to medial (bottom) axis showing a higher OMP-ir intensity in aAOB of an adult *O. degus* (A) and adult *O. lunatus* (B). C) Laboratory rats (*Rattus norvegicus*) exhibit a similar OMP-ir between both AOB portions. The site by which the vomeronasal nerve arrives at the AOB in each species is highlighted by an asterisk. Sections were counterstained with DAPI. Scale bar, 250 µm.

### 3.5 In degus, the bias to a larger aAOB emerges after birth and continues to increase with age

We investigated the development of the AOB and the establishment of an asymmetry as reported previously in adult degus (Suárez and Mpodozis 2009b). A volumetric study of the AOB was conducted in captivity-reared degus across different postnatal ages, from birth to adulthood. We measured the volumes of the overall AOB (considering the VNL, GL, EPL and M/TL) and each AOB portion at different postnatal ages: P1, P15, P30, P60, P150 and adults (P > 180). Similarly, the volume of the GL of the overall AOB and at each AOB portion was measured separately at each age.

#### Overall AOB and GL volumes

To determine whether AOB changes in size through the entire postnatal development, we measured the overall AOB volumes across different ages. In degus, we found that the AOB increased in overall size from P1 to adulthood (one-way ANOVA, F = 12.1, *p <* 0.001) (Table 1). The overall AOB increased 42% in area from P1 to P30 (*t* = 4.5, *p* < 0.001; Figure 7A). At P60, the AOB was 83% of its size at P30 (*t* = 1.6, *p* = 0.14). From P60, the AOB continued increasing, showing a maximum size in adults (36% larger than P60) (*t* = 3.4, *p* = 0.02; Figure 7A).

**Figure 7.**
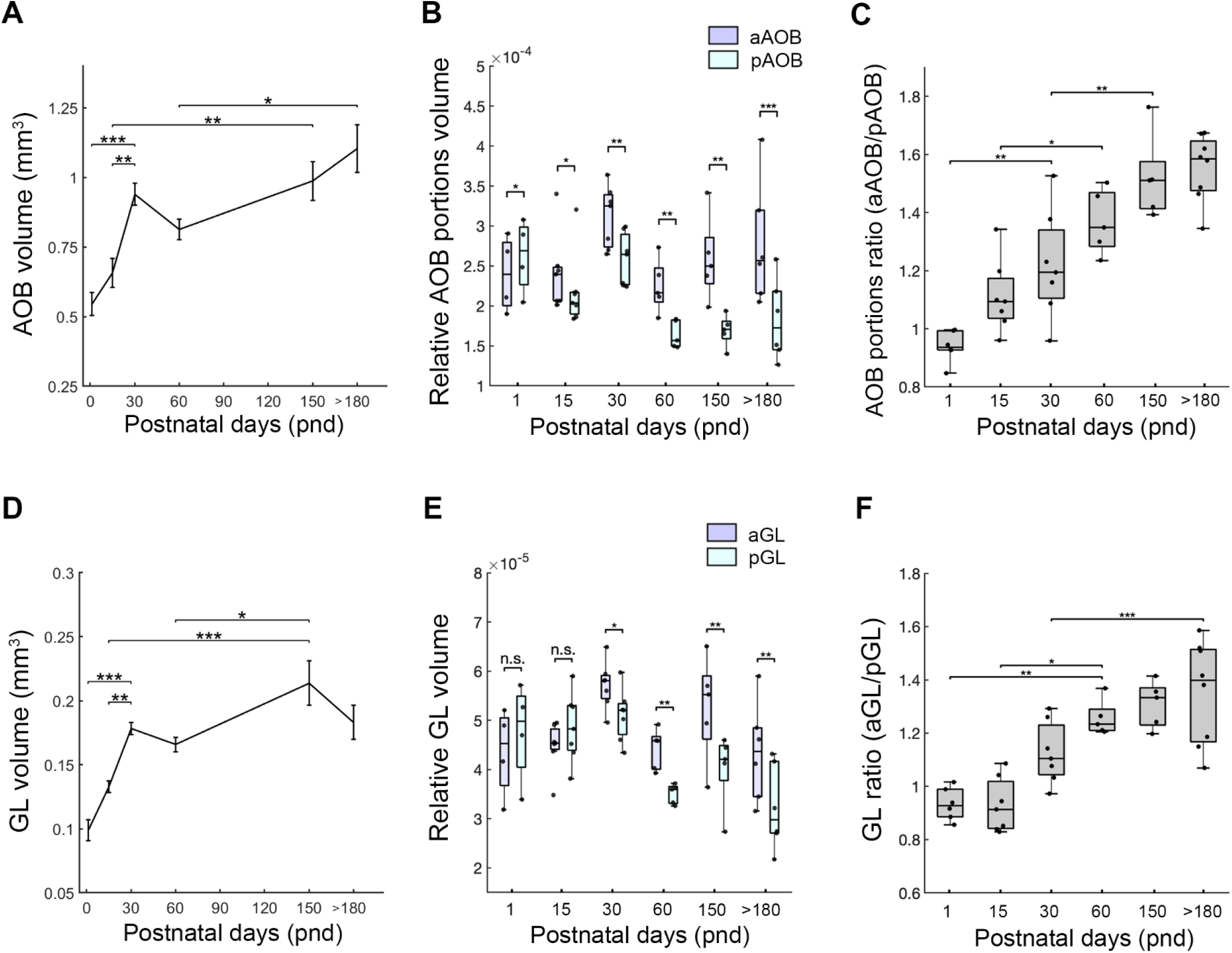
AOB morphometry and the establishment of the AOB asymmetry. A) Overall volume of the AOB measured at different postnatal ages: P1, P15, P30, P60, P150 and adults (P > 180). B) Boxplot graph showing the relative overall volume of aAOB and pAOB at different postnatal stages. A bias to a larger aAOB than pAOB volume is observed from P15. C) Boxplot graph showing the aAOB/pAOB overall size ratio. The adult-like bias extent is observed from P150. D) Overall volumes of the GL measured at different postnatal stages. A bias to a larger aGL than pGL volume is observed from P30. E) Boxplot graph showing the relative GL volumes of aGL and pGL in all the postnatal stages evaluated. A larger aGL than pGL is observed from P30. F) Boxplot graph showing the aGL/pGL size ratio. The maximum bias extent at GL is reached by adulthood. Significant differences between aAOB and pAOB measurements are shown with asterisks (n.s., non-significant, * p<0.05, ** p<0.01, *** p<0.001).

#### Overall volume of anterior and posterior AOB portions

To evaluate the time at which AOB asymmetry emerges and is established, we measured the normalized overall volume of the aAOB and pAOB separately, at different postnatal ages (Table 2). Unlike adult degus, which show a bias toward a larger aAOB than pAOB, newborn degus initially exhibit the opposite bias. At P1, pAOB was significantly larger than the aAOB (*t* = -4.3, *p =* 0.02; Figure 7B). By P15, the bias shifted to the typical adult pattern (aAOB > pAOB), with the aAOB being 11% larger than the pAOB (*t* = 2.6, *p =* 0.04). The larger size of aAOB than pAOB persisted at later ages: P30 (*t* = 3.2, *p =* 0.01), P60 (*t* = 6.6, *p =* 0.003), P150; (*t* = 5.7, *p =* 0.005). In adults, aAOB was 52% larger than the pAOB (*t* = 7.5, *p =* 0.001). From each age evaluated, we calculated the overall AOB volume ratio (aAOB/pAOB), which is a measurement of the extent of the AOB asymmetry. The AOB ratio increased progressively between P1 and P30 (*t* = 3.8, *p =* 0.004; Figure 7C) and again between P30 and P150 (*t* = 3.9, *p =* 0.004), indicating a growing extent of the AOB asymmetry across development. Thus, the bias toward a larger aAOB than pAOB emerges by the second first postnatal month in degus. The extent of this bias increases with age, reaching maximum values by adulthood. Thus, AOB asymmetry emerges within the first postnatal month, and increases in extent until adulthood.

#### GL volume in the anterior and posterior AOB portions

We measured the GL volume across different postnatal ages. Like AOB, GL exhibited an increase in size across ages (one-way ANOVA, F = 15.2, *p <* 0.001). GL increased 45% from P1 to P30 (*t* = 5.5, *p* < 0.001; Figure 7D). At P60, GL was 93% of its size at P30 (*t* = 0.9, *p* = 0.6) From P60, the GL volume continued to increase, showing a maximum size at P150 (22% larger than P60) (*t* = 3.3, *p* = 0.02). In adults, GL was 86% of its size at P150 (*t* = 2.2, *p* = 0.1). Thus, an increment of the AOB size and its GL occurs within the first month, with maximum sizes reached close to adulthood.

It was shown that degus exhibit a more and larger glomeruli at aAOB than pAOB (Suárez and Mpodozis 2009b). Similar to our analysis of the overall size of each AOB portion, we examined the normalized volumes of GL in the aAOB and pAOB (Table 2). At early ages, no significant differences in GL size were observed between aGL and pGL: P1 (*t* = -2.4, *p =* 0.1; Figure 7E) and P15 (*t* = -1.9, *p =* 0.1). However, at P30, the aGL was 12% larger than the pGL (*t* = 2.8, *p =* 0.03) and this bias persisted at P60 (*t* = 7.7, *p =* 0.002) and P150 (*t* = 6.1, *p =* 0.004). In adults, the aGL was 35% larger than the pGL (*t* = 4.4, *p =* 0.007). The GL ratio (aGL/pGL) showed a significant increase between P1 and P60 (*t* = 4.4, *p =* 0.001; Figure 7F) and between P30 and adults (*t* = 3.6, *p =* 0.01). Different to the overall size of each AOB portion, the bias towards a larger aGL than pGL in degus emerged later, at the end of the first postnatal month. The extent of this AOB asymmetry continued to increase with age, reaching a maximum extent slightly before adulthood. Thus, AOB asymmetry emerges within the first postnatal month, and increases in extent until adulthood.

### 3.6 In degus, the AOB asymmetry results from a smaller increase of pAOB compared to aAOB across postnatal development

We examined how changes in the sizes of aAOB or pAOB across ages contribute to the development of the AOB asymmetry. Linear regression analysis revealed that the relative volumes of pAOB and pGL decreased with age compared to the aAOB (ANCOVA: F = 6.92, *p =* 0.01; Figure 8A) and aGL (ANCOVA: F = 6.65, *p =* 0.01; Figure 8B), respectively. During this period, brain volume increased progressively (Figure 8C). Overall, the results indicate that while the AOB portions increase with brain size, the pAOB and its corresponding GL show a comparatively smaller increase than aAOB during the same period, resulting in an increase in the extent of the AOB asymmetry throughout postnatal development.

**Figure 8.**
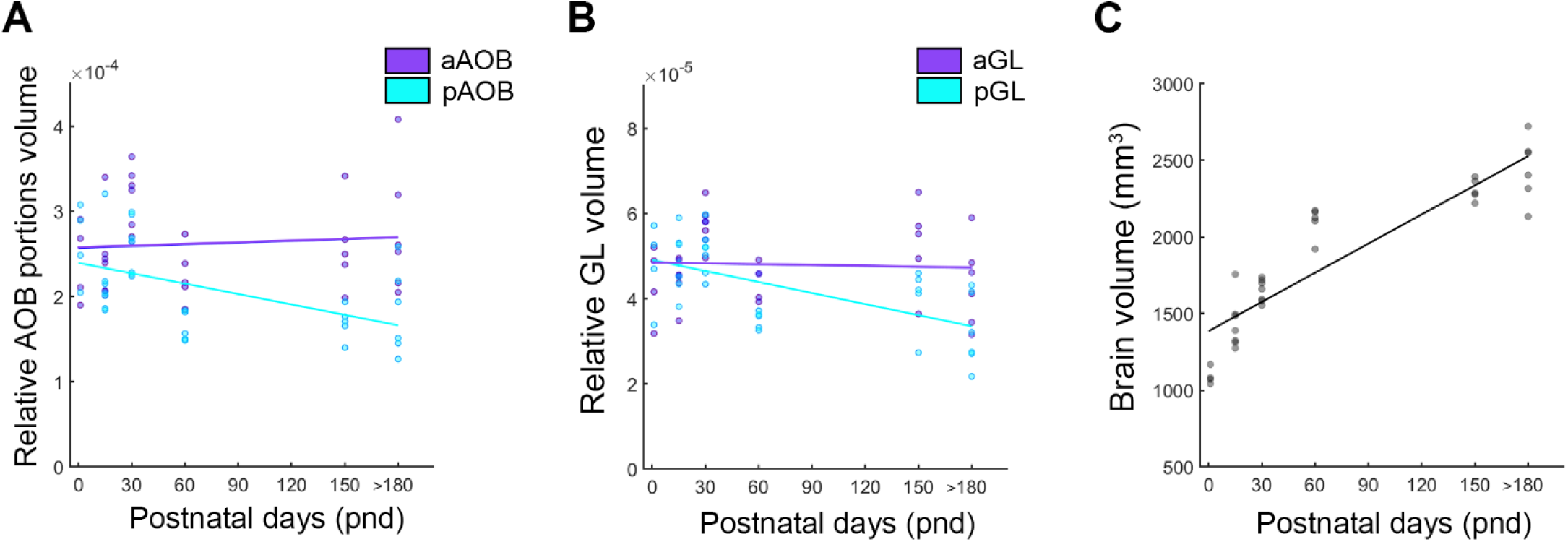
pAOB shows a minor growth rate than aAOB acros postnatal ages. A) Linear regression plot over the relative AOB portions (aAOB and pAOB) volumes, obtained at P1, P15, P30, P60, P150 and adults (P > 180). aAOB and pAOB linear functions show a significant difference in their slopes (ANCOVA: F = 6.92, *p* = 0.01). B) Linear regression plot over the relative GL volumes of each portion. aGL and pGL linear functions show a significant difference in their slopes (ANCOVA: F = 6.65, *p* = 0.01). C) Linear regression plot on brain volumes obtained at all the postnatal ages evaluated.

### 3.7 The extent of AOB asymmetry varies among *O. degus* raised in different ecological contexts

To explore how ecological and social rearing contexts influence AOB asymmetry, we compared the volumetric measurements of the aAOB and pAOB in adult degus reared in captivity against two other conditions: field-captured and isolation-reared.

#### AOB morphometry of field-captured and isolation-reared degus

In our study, field-captured degus exhibited a prominent bias toward a larger aAOB, consistent with a/our previous report (Suárez and Mpodozis 2009b). The overall volume of the aAOB was 69% larger than pAOB (*t* = 7.3, *p =* 0.001, *n* = 6; Figure 9A). Similarly, the volume of the GL showed the same bias, with the aGL being 62% larger than the pGL (*t* = 7, *p =* 0.001, *n* = 6; Figure 9B). In contrast, degus reared in isolation did not display a significant difference in size between aAOB and pAOB, neither in overall AOB volume (*t* = 2.5, *p* = 0.1, *n* = 3) nor in the GL (*t* = 1.8, *p* = 0.2, *n* = 3).

**Figure 9.**
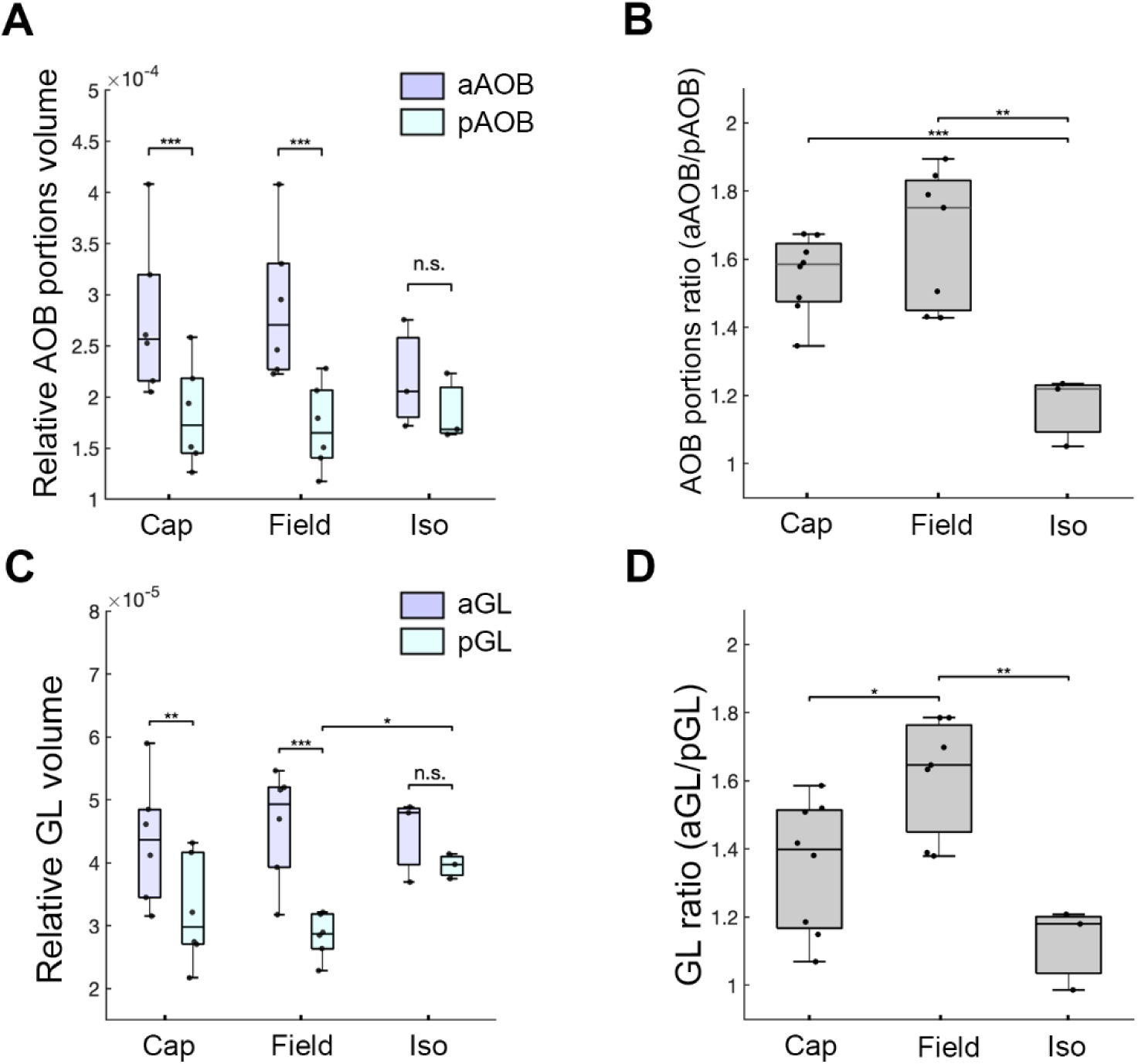
*Octodon degus* raised in different social contexts show differences in the extent of the AOB asymmetry. A) Boxplot graph showing the relative overall volume of AOB portions measured in captivity-reared (Cap), field-captured (Field) and isolation-reared (Iso) individuals. B) Boxplot graph showing the aAOB/pAOB overall size ratio. C) Boxplot graph showing relative GL volumes meaured in captivity-reared (Cap), field-captured (Field) and isolation-reared (Iso) degus. D) Boxplot graph showing the aGL/pGL size ratio of captivity-raised, field-raised and isolation-raised individuals. Significant differences are indicated with asterisks (* p<0.05, ** p<0.01, *** p<0.001).

#### Comparison between degus raised in different social contexts

To determine if social context in which degus raise influences on the AOB shaping, we first compared the volumes of the AOB portions and their GLs volumes between field-captured and captivity-reared individuals. No significant differences in overall AOB size were observed between field-captured and captivity-reared degus for aAOB (*t* = -0.3, *p* = 0.8, *n_field_* = 6, *n_cap_* =6; Figure 9A) or pAOB (*t* = 0.4, *p* = 0.7, *n_field_* = 6, *n_cap_* =6). There was no significant difference in the overall AOB ratio (aAOB/pAOB) between captivity-reared and field-captured degus (*t* = -1.4, *p* = 0.2, *n_field_* = 7, *n_cap_* =8; Figure 9B). Similarly, in the GL, no differences were found at the aGL (*t* = -0.4, *n_field_* = 6, *n_cap_* =6; Figure 9C) or pGL (*t* = 1, *p* = 0.3, *n_field_* = 6, *n_cap_* =6) between field-captured and captivity-reared degus. The GL volume ratio (aGL/pGL) of field- captured degus was significantly bigger in comparison to captivity-reared degus (*t* = -2.8, *p* = 0.02, *n_field_* = 7, *n_cap_* =8; Figure 9D). Thus, in captivity conditions, degus developed a less asymmetric GL compared to degus raised in the wild.

Compared to field-captured animals, isolation-reared degus did not differ significantly in overall size of the aAOB (*t* = -1.5, *p* = 0.2, *n_field_* = 6, *n_iso_* =3; Figure 9A) or pAOB (*t* = -0.5, *p* = 0.6, *n* = 3). However, isolation-reared degus exhibited a significantly smaller overall AOB ratio compared to field-captured degus (*t* = 3.9, *p* = 0.004, *n_field_* = 7, *n_iso_* =3; Figure 9B). At the GL, isolation-reared degus showed a significantly larger pGL compared to field-captured animals (*t* = -5, *p* = 0.002, *n_field_* = 6, *n_iso_* =3; Figure 9C), whereas no differences were observed in the aGL (*t* = 0.2, *p* = 0.8, *n_field_* = 6, *n_iso_* =3). The GL volume ratio (aGL/pGL) of isolation-reared degus was significantly smaller in comparison to field-captured (*t* = 4.5, *p* = 0.002, *n_field_* = 7, *n_iso_* =3; Figure 9D). Thus, AOB morphometry in isolation-reared degus is highly distinct from that observed in degus reared in the wild.

Compared to captivity-reared animals, isolation-reared degus did not differ significantly in the overall size of the aAOB (*t* = 1.2, *p* = 0.3, *n_cap_* = 6, *n_iso_* =3; Figure 9A) or pAOB (*t* = -0.1, *p* = 0.9, *n_cap_* = 6, *n_iso_* =3). The AOB ratio, however, differed significantly between these groups (*t* = 5.1, *p =* 0.001, *n_cap_* = 8, *n_iso_* =3; Figure 9B). At the GL, no differences between groups were found between both aGL (*t* = -0.2, *p* = 0.9, *n_cap_* = 6, *n_iso_* =3; Figure 9C) and pGL (*t* = -1.4, *p* = 0.2, *n_cap_* = 6, *n_iso_* =3). Also, no differences were observed in GL ratio between captivity-reared and isolation-reared degus (*t* = 1.9, *p* = 0.09, *n_cap_* = 8, *n_iso_* =3; Figure 9D). Thus, captivity-reared degus exhibit an AOB morphometry which is closer to that observed in isolation-reared degus.

Overall, our results suggest that environmental context influences the development of AOB asymmetries, with natural rearing promoting the typical bias towards a larger aAOB than pAOB.

## 4. Discussion

Our results show that the asymmetry previously reported in adult degus in AOB (Suárez and Mpodozis 2009b) emerges within the first postnatal month. Interestingly, in newborns, the AOB partitioning is biased toward pAOB, opposite to adults. This bias shifts towards an aAOB that is larger than pAOB between the second and fourth postnatal week, and it increases with age, reaching a maximum extent by adulthood. The AOB asymmetry mainly results from a minor growth of pAOB compared to aAOB across development. Both the VNO and AOB exhibit anatomical and molecular changes throughout the first postnatal month. Notably, from the second week on, V1R neurons showed a stronger OMP expression than V2R neurons in VNS structures. At the end of the first month the GL exhibited a larger overall size and more well-defined glomeruli surrounded by interneurons, closely resembling adult GL anatomy. We also showed the influence of environmental factors on AOB asymmetry; it is less pronounced in individuals reared in captivity and absent in animals reared in isolation. The more prominent AOB asymmetry develops mainly in wild degus.

### 4.1 The maturation of AOB circuitry

Differences in the time of maturation of MOS and VNS have been reported among rodents. MOS structures exhibit high level of maturation in newborns of different mammalian species such as marsupials (Shapiro et al. 1997) and rodents (Salazar et al. 2006; Torres et al. 2020), suggesting a functional role from early postnatal ages. In newborn precocial rodents such as capybaras, which born with open eyes, AOB exhibits an advanced pattern of maturity in both anatomy and gene expression (Torres et al. 2020), suggesting a functional role for the VNS right after birth. In contrast, altricial mice and rats open their eyes two weeks after birth and remain fully dependent on maternal care during this period. Different to capybaras, mice exhibit a stratified AOB with glomeruli by postnatal day 5 (Salazar et al. 2006), and important changes in glomeruli formation are observed during the first month (Hovis et al. 2012). In altricial species, a functional maturation of the VNS is suggested to occur within the first month, based on anatomy (Salazar et al. 2006; Hovis et al. 2012), gene expression of key molecular markers (Salazar et al. 2006), and local activity pattern at AOB (Sugai et al. 2005).

In contrast to mice, degu pups are born with open eyes, and they exhibit scavenging behavior during their first week (Veloso 1997; Reynolds and Wright 1979). At birth, MOS structures exhibit substantial maturation, suggesting a functional role of MOS at early postnatal. Newborns possess a well stratified AOB, although glomeruli are scarce. By the second week, AOB still exhibits important immature traits, with a small overall size and a low incorporation of interneurons in the AOB circuitry compared to posterior ages, reflecting an ongoing maturation. The expression of key molecular VNS markers as OMP, and indicator of anatomical maturation of olfactory neurons, is first observed by the second postnatal week. Our results suggest that the maturation of VNS in degus is established during the first postnatal month, resembling what is observed in altricial species.

In degus, a low presence of interneurons in the GL was observed during the first two postnatal weeks, compared to later developmental stages. Interneurons are crucial for modulating local olfactory bulb activity and coding odor information (Lledo et al. 2008; Maksimova et al. 2019). Through dendrodendritic synapses, they provide feedback and lateral inhibition and synchronize mitral and tufted neurons’ outputs, influencing signals reaching telencephalic targets (Aungst et al. 2003; McGann et al. 2005; Huang et al. 2013). In mammals, prolonged neurogenesis allows the incorporation of new interneurons into the olfactory bulb from early ages to adulthood, conferring high plasticity to olfactory circuits across development (Alvarez-Buylla and García-Verdugo 2002; Rochefort et al. 2002; Rochefort and Lledo 2005; Lledo et al. 2006; Portillo et al. 2012). In degus, the low presence of interneurons in GL at early ages likely reflects differences between juveniles and adults in neuronal modulation of the AOB local activity.

In rodents, the maturation of local AOB activity is established after birth. In rats, the local AOB activity exhibits adult-like patterns by P18 (Sugai et al. 2005). It coincides with the moment at which AOB reaches maximum size (Rosselli-Austin et al. 1987), followed by a decrease in size until it reaches adult-like size by P60. AOB in degus exhibits an increase in size during the first postnatal month. At the end of the first month, AOB possesses abundant, well-defined glomeruli surrounded by interneurons. Our findings suggest that in degus, a possible transition to a more mature AOB, in terms of the modulation of the AOB local activity occurs by the end of the first postnatal month. Future experiments on the neuronal activity of AOB across early development, searching for an “electrophysiological maturation” time of the AOB circuitry, as suggested in rats (Sugai et al. 2005) may help to clarify this point.

At early postnatal ages, the establishment of connections at AOB is dependent on activity triggered by stimuli (Hovis et al. 2012). Based on our results, we suggest that the VNS activity in degus at early ages may influence the maturation of the AOB circuitry. Olfactory cues learned at early ages can influence olfactory-guided behaviors in adult degus (Márquez et al. 2015; Márquez et al. 2025), suggesting that an effect of early experience on the olfactory circuitry of degus, including VNS, may occur.

One of our surprising findings was the stronger OMP expression observed in apical VNO neurons and their projections to the AOB from P15 onwards, compared to basal VNO neurons, found in degus and lunatus. To our knowledge, no reports have documented differences in OMP expression between vomeronasal populations within eutherians. Similar patterns were previously reported in the opossum *Monodelphis domestica* (Marsupialia, Didelphidae), in which more intense OMP-ir is seen in apical VNO neurons compared to basal neurons from weaning (P60; Shapiro et al. 1997). That difference in OMP expression is well established at the time animals engage in more complex social interactions, suggesting an association between the cellular maturation of VNS and behavior (Shapiro et al. 1997).

In rodents such as mice and rats, OMP expression at VNS appears early, but reaches adult-like levels by the end of the first month (Farbman and Margolis 1980). In capybaras, however, OMP expression in the VNS is well established by birth (Torres et al. 2020). In degus, OMP is expressed by all vomeronasal neurons at birth; however, a more mature OMP pattern appears by the second week. This indicates OMP experience a prolonged period of maturation through the first postnatal month in degus, resembling that of altricial species.

In rodents, OMP expression in the main olfactory system influences particular response properties of olfactory neurons such as response speed and selectivity and sensitivity to odorants (Buiakova et al. 1996; Kass et al. 2013). It is possible that differences in OMP expressions in octodontids and *M. domestica* may translate into differences in pheromone processing between vomeronasal pathways, supporting the notion that these participate in different behavioral modalities (Suárez and Mpodozis 2009b).

Overall, we show that degu VNS exhibits remarkably immature traits at birth. The maturation of the AOB is reflected in the formation of glomeruli surrounded by interneurons and in differential expression of OMP between vomeronasal pathways. We suggest that a highly mature VNS, as observed in adult degus, is established by the end of the first postnatal month.

### 4.2 The development of asymmetric AOB in degus

One of the most remarkable traits in degus is the asymmetry of the AOB, with the aAOB being larger and containing more and larger glomeruli than pAOB (Suárez and Mpodozis 2009b). How could the AOB asymmetry be established across ontogeny? Our findings show that in degus, this bias emerges within the first month and exhibits an increase in extent over later ages.

In 2-week-old degus, the aAOB already shows a larger volume than the pAOB. We did not count the number of V1R and V2R inputs reaching the AOB at each age evaluated. However, our results on OMP expression at AOB suggest that a greater number of V1R neurons expressing stronger OMP projecting to the aAOB than basal neurons projecting to pAOB. Interestingly, a larger number of V1R neurons projecting to aAOB at P15 does not result in a larger aGL than pGL during the second postnatal week. It may indicate that at early postnatal ages, more densely packed connections are established in aAOB compared to pAOB.

From the end of the first month, the increase in the size of each AOB portion occurs, with the pGL showing a slower growth rate than aGL, resulting in an asymmetric AOB. In degus, changes in glomeruli size are thought to occur over this period, resulting in more and larger glomeruli in the aAOB than in pAOB by adulthood (Suárez and Mpodozis 2009b). Olfactory systems, including VNS, exhibit plasticity in the number and size of glomeruli across postnatal development. Changes in the number of olfactory neurons and the size of the glomeruli have been reported in adult mice exposed to specific stimuli under conditioning tests (Kerr and Belluscio 2006; Jones et al. 2008). Rapid remodeling of olfactory neuron inputs into the OB driving by activity contributes to the OB plasticity (Cheetham et al. 2016). The remodeling of synaptic connectivity across later ages in VNS has been associated with adult neurogenesis, which is adjusted to environmental changes (for review see Oboti and Peretto 2014). In degus, the AOB asymmetry may result from changes in the size and/or in the number of glomeruli over later ages, driven by experience.

Along with the incorporation of new vomeronasal neurons in degus AOB, we found that mitral and tufted neurons exhibit changes in their distribution with age. Activity dependent synaptic remodeling of the mitral and tufted neurons dendrites occurs during the first postnatal month (Hovis et al. 2012). Postnatally, the number of mitral and tufted projection neurons can be influenced by hormonal changes (Valencia et al. 1986). In rats, changes in size of AOB layers, including the M/TL, follow changes in the overall AOB size across postnatal ages, with the exception of GrL (Rosselli-Austin et al. 1987). Thus, the number and distribution of secondary projection neurons in the AOB fluctuates with age and experience. In adult degus, we reported previously that aAOB possesses more mitral and tufted neurons than pAOB (Fernández-Aburto et al. 2019), but how and when such a bias is established is unknown. The study of development of mitral tufted neurons in degus resulting in their asymmetric distribution in the AOB may better clarify this point.

Overall, the combination of increasing neuron numbers and activity-dependent remodeling likely underpin the emergence and establishment of the AOB asymmetry observed in adult degus.

### 4.3 Differences in the AOB asymmetry among octodontids raised in different social contexts

In mammals, social interactions have been suggested to play a significant role shaping the AOB. Early exposure to environmental odors affects social behavior later in life (Broad and Keverne 2012). In mice, male-male social interactions led to changes in the excitatory/inhibitory synaptic balance between mitral and granular AOB neurons (Cansler et al. 2017). In hamsters, the size of synaptic contact zones and the somata area of M/T neurons in the AOB become larger in animals raised in a more socially enriched context (Matsuoka et al. 1994, 1998). In the common shrew *Tupaia glis* (Scadentia), the total AOB volume was found to be larger in individuals raised in the field in comparison with those raised in captivity (Frahm et al. 1984). Recently, we reported that *Octodon lunatus*, a degu sister species exhibiting less social interaction (Sobrero et al. 2014), exhibits a diminished AOB bias in comparison to degus (Fernández-Aburto et al. 2019).

Our findings show that the development of AOB asymmetry is strongly influenced by social experience. In degus, scent marking is exuberant, involving urine and dustbathing. Dustbathing is performed at sites that are frequently urine marked (Wilson and Kleiman 1974). Agonistic encounters in degus may also include dustbathing (Fulk 1976). In captive degus, males decrease the urine marking when exposed to scent marks of conspecifics of the same sex (Kleiman 1975). In addition, degus are capable of discriminating between socially familiar and unfamiliar scents deposited by dustbathing (Ebensperger and Caiozzi 2002). Thus, behaviors involving the use of pheromone cues in degus are diverse and influenced by conspecifics.

Different from behavior in the field, in restricted social contexts as observed in laboratory conditions, a reduction in the diversity of social interactions involving pheromone signaling compared to wild degus is expected to occur. We observed that degus raised in captivity or in controlled isolation conditions exhibit a less aAOB biased or even unbiased AOB, especially in the GL. In the laboratory, degus were reared in smaller and less diverse groups and were confined to a more restricted space than in the field. We suggest that in such conditions, most of the diversity in social interactions seen in wild is reduced, including those in which volatile cues are preferentially used. In restricted conditions, pheromonal communication in degu encounters is expected to occur with no restrictions through development, but in a different way than expected in wild, social contexts. It could lead to the unbiased use of both volatile and non-volatile cues, promoting the equal increase in size of both aAOB and pAOB.

In comparison to wild degus, aGL size in individuals raised in restricted conditions exhibit a similar size. In contrast, pGL increases in size in animals reared in isolation conditions, equaling aGL. We reported here that the development of AOB asymmetry results from a minor growing rate of pAOB traits compared to aAOB. Overall, we suggest that in degus, the development of the pAOB traits, as GL size, is more labile than in aAOB to changes in social context.

The large, less size-biased GL observed in degus reared in restricted social conditions is different compared to the less size-biased AOB reported in *O. lunatus* (Fernández-Aburto et al. 2019). In comparison to field-reared degus, lunatus exhibits a significantly smaller aGL, having a similar in size as the pGL. Lunatus exhibits a lesser extent of sociality, with a reduced number of encounters during the day compared to degus (Sobrero et al. 2014). Under such conditions, *O. lunatus* are expected to get involved in less diverse and less frequent encounters during life in comparison to wild degus. This could lead to a decreased reliance on pheromonal communication, and thus, to the development of a small, less biased AOB in *O. lunatus* than degus. We suggest that different to degus, in which rearing in restricted social conditions affect the pAOB development more, in *O. lunatus* a lesser extent of sociality would affect the aAOB development more. Thus, the AOB shape and compartmental organization among octodontids is affected differently depending on the social context.

Overall, our results support the hypothesis that greater social complexity and pheromonal communication in wild degus drive the enhancement of aAOB size and the smaller increase of pAOB size compared to aAOB through postnatal development. It highlights the plasticity of the AOB shape in response to social experience, emphasizing the importance of social context in neural development and the establishment of sensory specializations.

### 4.4 Conclusions

During postnatal development, *O. degus* gradually develop a bias toward a larger aAOB starting by the second postnatal week. This bias increases with age and is mainly driven by the minor growing rate of pAOB traits, particularly in the GL, leading to a smaller pAOB in adults. In environments with reduced social interactions, such as captivity and isolation conditions, the AOB asymmetry decreases in extent as pAOB traits increase in size. Overall, our results support the hypothesis that the emergence and establishment of the AOB asymmetry in degus is driven by social experience. During the time when VNS traits become more mature, experience-dependent regulation in the development of the AOB asymmetry through ontogeny is observed.

## Author contributions

PFA conceived and performed the experiments, analyzed the results and wrote the manuscript. SED, KB, JMU and RS collected the material, performed experiments and analyzed the results. JMP conceived the experiments, supervised the project and co-wrote the manuscript.

## Data availability statement

The data and images that support the findings of this study are available from the corresponding authors upon reasonable request.

## Acknowledgements

The authors gratefully acknowledge Dr. Luis Ebensperger, Dr. Pablo Sabat and Dr. Rodrigo Vasquez for providing us with animals. Thanks to Solano Henriquez, Elisa Sentis and Tatiana Guarnieri for their technical assistance. We thank Dr. Sarah Pallas for her valuable comments on the manuscript. This work was supported by MECESUP Doctoral Fellowship UCH0713, FONDECYT 1170027 and FONDECYT 3150306. The authors declare no competing financial interests.

## Notes

### Competing Interest Statement

We would like to note that Dr. Jorge Mpodozis is an Editorial Board member of this submitted journal and one of the corresponding authors of this article. To minimize bias, he must to be excluded from all editorial decision making related to the acceptance of this article for publication. The authors declare no other conflict of interest.

